# The geometric determinants of programmed antibody migration and binding on multi-antigen substrates

**DOI:** 10.1101/2020.10.12.336164

**Authors:** Ian T. Hoffecker, Alan Shaw, Viktoria Sorokina, Ioanna Smyrlaki, Björn Högberg

## Abstract

Viruses and bacteria commonly exhibit spatial repetition of surface molecules that directly interface with the host immune system. However the complex interaction of patterned surfaces with multivalent immune molecules such as immunoglobulins and B-cell receptors is poorly understood, and standard characterization typically emphasizes the monovalent affinity. We developed a mechanistic model of multivalent antibody-antigen interactions as well as a pipeline for constructing such models from a minimal dataset of patterned surface plasmon resonance experiments in which antigen pattern geometries are precisely defined using DNA origami nanostructures. We modeled the change in binding enhancement due to multivalence and spatial tolerance,i.e. the strain-dependent interconversion of bound antibodies from monovalently bound to bivalently bound states at varying antigen separation distances. The parameterized model enables mechanistic *post hoc* characterization of binding behavior in patterned surface plasmon resonance experiments as well as *de novo* simulation of transient dynamics and equilibrium properties of arbitrary pattern geometries. Simulation on lattices shows that antigen spacing is a spatial control parameter that can be tuned to determine antibody residence time and migration speed. We found that gradients in antigen spacing are predicted to drive persistent, directed antibody migration toward favorable spacing. These results indicate that antigen pattern geometry can influence antibody interactions, a phenomenon that could be significant during the coevolution of pathogens and immunity in processes like pathogen neutralization or affinity maturation.

## 1 Introduction

Due to their multiple binding domains, immunoglobulin molecules like the bivalent IgG antibody exhibit complex interactions with multivalent antigens, i.e. clusters of multiple copies of molecules or molecular domains occurring on the order of 1-30 nm separation distances. Multivalent interactions enhance the stability of binding interactions by enabling simultaneous attachment of multiple ligands, increasing the statistical ratio of bound to unbound states and extending the residence times of bound antibodies[9, 8].

Many pathogenic surfaces exhibit spatial repetition at length scales relevant to antibody multivalence. Viral capsid proteins undergo self-assembly into periodic patterns [15], and some neutralizing antibodies achieve their high affinity and neutralization capability through bivalence [3, 14]. Self-assembling crystalline arrays of surface-layer (S-layer) proteins, the outermost structure on many bacteria and archaea, are a major contact point between pathogen and host [1] and are implicated as mediators of innate [13] and adaptive immunity [6, 5]. Their repetitive organization may be integral to their immunological role, as their removal from cell surfaces was seen to reduce immune response [12]. Multivalence is also likely an important factor during affinity maturation of antibodies and thus in vaccine design [16, 7].

Antibody interaction with patterned surfaces presents a challenge for mathematical modeling as it is a many-bodied problem occurring on timescales of seconds to minutes. Such systems are too computationally expensive for full-atom molecular simulation. Models treating antibodies and antigens as abstract binding and non-binding units have been the most successful at capturing relevant dynamics, and have historically treated multivalence as a function of ligand coating density whereby multivalence emerges as ligand nearest-neighbor distance statistically decreases with higher densities [10]. More recently, coarse-grained molecular simulations have been used fruitfully to investigate the role of antibody multivalence on binding kinetics [2]. However a challenge of precisely calibrating such models remains due to the absence of both experimental tools to independently assess monovalent and multivalent binding dynamics as well as a computational pipeline to connect such data to models.

The patterned surface plasmon resonance (PSPR) technique (Fig 1 a) enables measurement of binding kinetics on precise, monodisperse patterns of ligands, achieving robust control of geometry through the use of DNA origami nanostructures [11]. Herein, we demonstrate a pipeline for the automated conversion of PSPR data into a flexible, experimentally parameterized model of antibody interaction with arbitrarily complex multivalent surfaces. The model is based on a coarse-grained simplification of bivalent antibody binding to antigens as a discrete Markov process with distinct states: empty antigen, monovalent antibody-antigen complexes, and bivalent antibody-antigen complexes with transitions between these states governed by elementary rates (Fig 1 b). From this basis, the dynamics of more complex patterns of multiple antigens can be reduced (Fig 1 c) to combinations of these elementary states. We show how the spatial tolerance, i.e. the range and impact of antigen separation distances on bivalent binding kinetics (Fig 1 d), can be exploited in the design of antigen patterns to modulate the effective binding affinity, the walking speed of antibody migration, and even the direction of antibody migration on patterned surfaces. A causal linkage between pattern geometry and antibody dynamics could play a role in the co-evolution of adaptive immunity and pathogens.

**Fig. 1.**
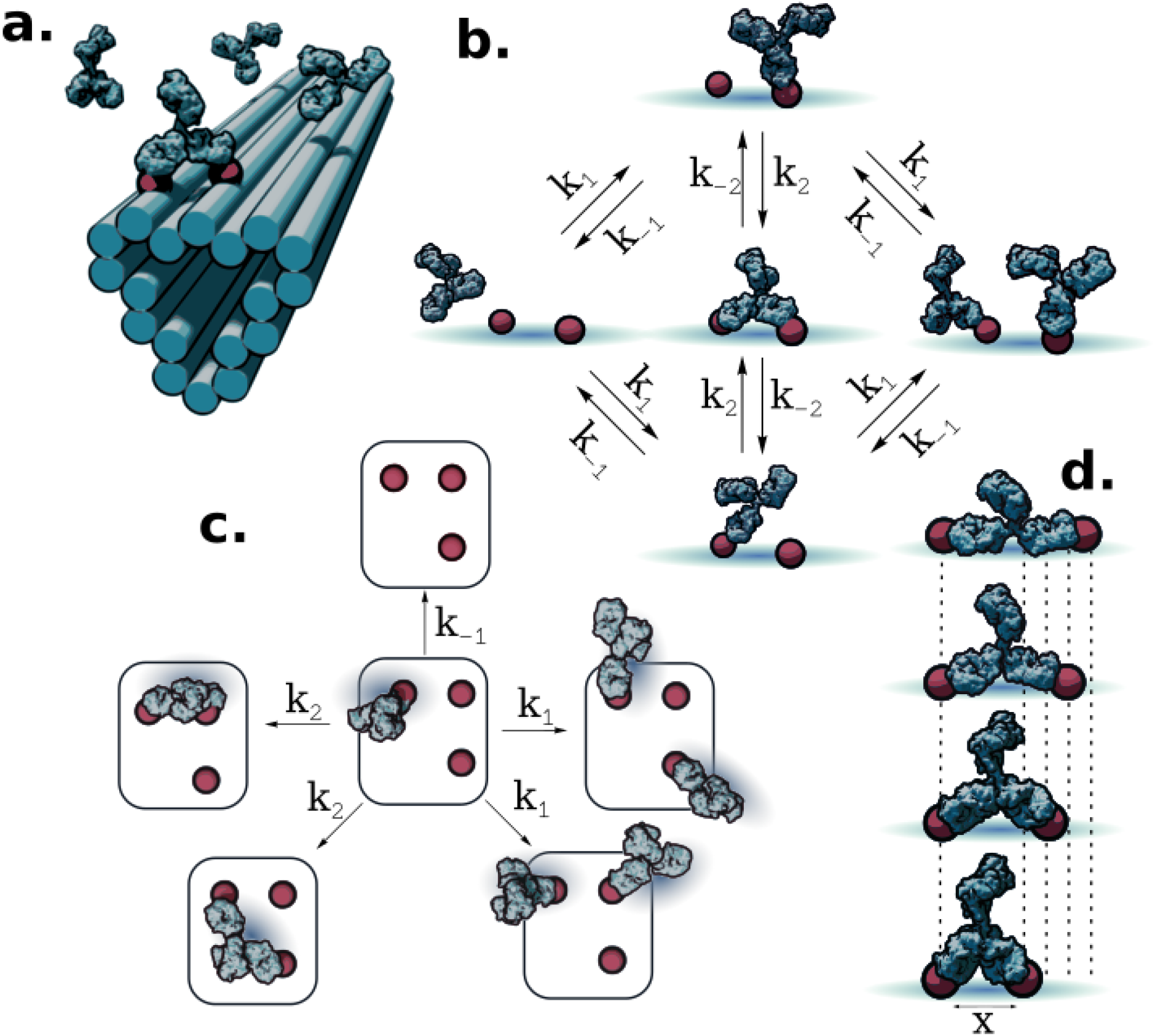
Scheme for modeling binding dynamics of antibodies on multi-antigen substrates. a: Illustration of patterning concept, whereby small molecule antigens (haptens) are arranged using short, flexible tethers at well-defined locations on DNA origami nanostructures. This enables multivalent interaction of antibodies with antigen patterns. b: Markov model of antibody binding whereby only basic binding/unbinding and bivalent interconversion processes are used to couple discrete monovalent and bivalent binding states. c: Model extension to more complex pattern geometries is accomplished by separating the system into elementary transitions between states comprising different combinations of empty and monovalently or bivalently occupied antigens. d: Pairs of antigens separated by different lengths elicit differing antibody binding kinetics due to the separation distance dependent impact of antibody structure on the chance of bivalent interconversion.

## 2 Results

We developed a model parameterization pipeline based on progressive fitting of the transient SPR profiles for first monovalent and then bivalent binding processes in order to reduce degrees of freedom at each stage of fitting. In the first 1-antigen configuration stage, we used an anti-digoxygenin single-cycle-kinetics program whereby progressively higher concentrations of antibody were exposed to the immobilized antigen substrate (Fig 2 a). This program was performed with a 1-antigen configuration (Fig 2 b, d) in order to parameterize our Markov model yielding respective association and dissociation rates *k*_1_ = 2.1 × 10^7^*M*^−1^*s*^−1^ and *k*_−1_ = 5.3 × 10^−4^*s*^−1^ as well as a monovalent dissociation constant *K*_*D*1_ = 2.5 × 10^−11^ *M*. Parameterizating intercoversion between monovalent and bivalent states was done by fixing the previously determined monovalent parameters and fitting the model to experiments involving multiple adjacent antigens via adjustment of *K*_*D*2_ or the interconversion constant defined by the ratio of reverse and forward interconversion rates. For structures configured with 2 antigens separated by ≈ 14 *nm*, we find *K*_*D*2_ = 5.8 × 10^−2^ (Fig 2 c, e).

**Fig. 2.**
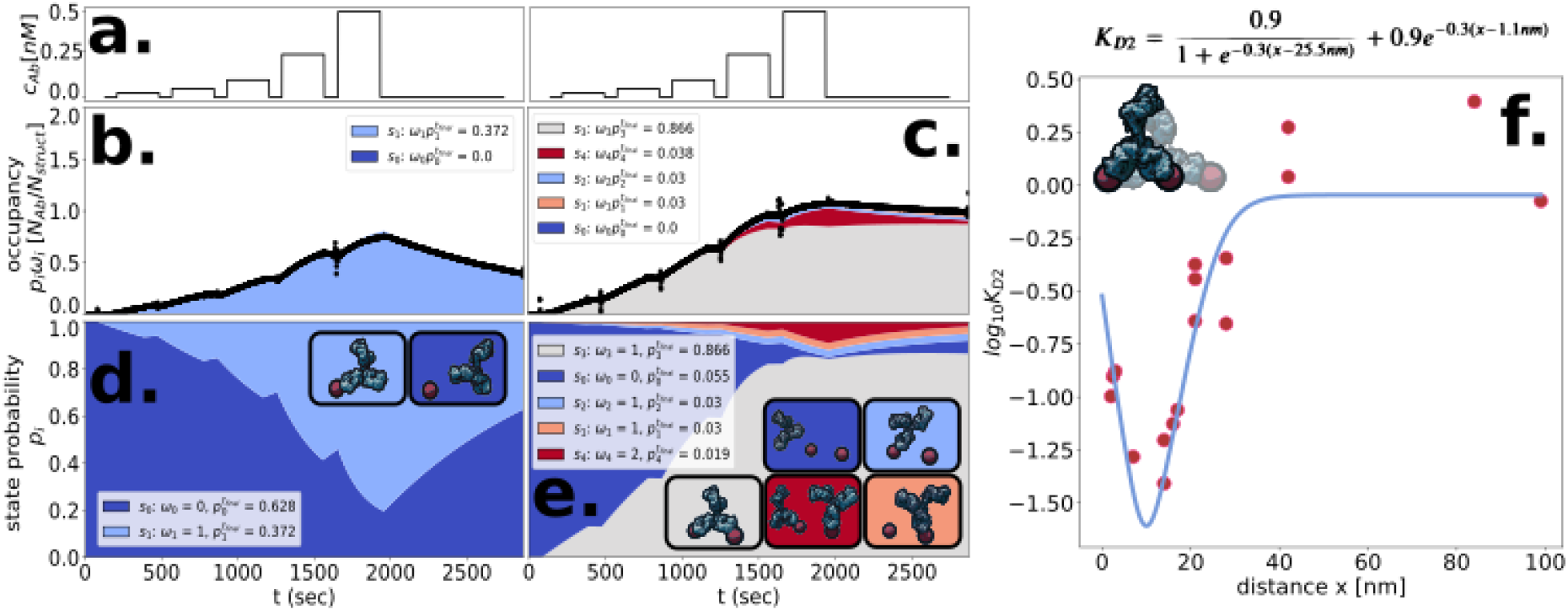
A progressive fitting pipeline for obtaining a parameterized model from minimal experimental data. a: Concentration versus time plot of antibody solution exposed to patterned antigen substrates in a single cycle kinetics PSPR experiment. b and c: Experimental binding kinetics data (black line) of 1 antigen and 2 antigen configurations respectively, superimposed over the occupancy calculated from the parameterized model. Model occupancy is divided and colored according to state, with height corresponding to the state’s contribution to total antibody occupancy per structure. d and e: 1 and 2 antigen configuration respective transient state probabilities stratified according to model prediction, colored and stacked to satisfy the normalization condition whereby all probabilities add to 1. f: Interconversion constants (magenta points) plotted versus 2-antigen configuration separation distance *x* and the fitted spatial tolerance model (blue line).

Applying progressive fitting to PSPR runs with structures patterned with 2 adjacent antigens of varied separation distances, we found the internal conversion process to vary accordingly. Small and large separation distances correspond to reduced bivalence, i.e. larger *K*_*D*2_. We constructed a phenomenological equation modeling the interconversion constant (Fig 2 f). The model is composed of a logistic tension-term representing the reduced bivalence at large separation distances and an exponential compression-term representing the penalty to bivalence observed at extremely close separation distances. The interconversion constant is thus a function of adjacent antigen separation distance with the form

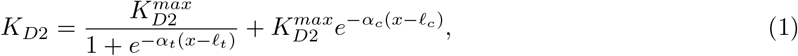

where *ℓ*_*t*_ and *ℓ*_*c*_ are characteristic lengths defining the scale of the tensile and compressive terms, *α*_*t*_ is the sharpness of the tensile penalty, *α*_*c*_ is the decay parameter of compressive penality to bivalence, and 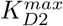 is the value of *K*_*D*2_ at which contributions of bivalence to binding dynamics is vanishingly small.

In order to determine the dependence of system bivalence on solution phase concentration, we used the parameterized model to obtain the steady state probability distributions for a range of solution phase concentrations (Fig 3 a). This revealed concentration regimes of differing dominant states: empty, bivalent, and saturated monovalent at respectively low, medium, and high solution phase concentrations. Entropic maxima occur at transitions between these domains (Fig 3 b), and transitions in bivalent and monovalent contributions to chemical potential occur in accordance with the transition from bivalent to saturated monovalent regimes (Fig 3 c). In addition to simulating de novo steady state properties, the model enables us to simulate transient dynamics of hypothetical systems with arbitrary geometries and arbitrary timing in the introduction of different solution phase concentrations. To demonstrate this, we simulated the evolution of state probability for a system with antigens arrayed in a 12 × 16 × 20 nm right triangle (Fig 3 d) with arbitrarily timed introduction of solution phase concentrations: 5 × 10^−4^, 6 × 10^−2^, 5 × 10^−1^, 1 nM at 0, 1000, 2000, and 3000 seconds respectively. This allowed us to track the evolution of entropy (Fig 3 e) as the system approaches steady state, revealing differences in both the total and individual values among states under the influence of different solution concentrations and time evolution. We also saw the time evolution of the number of antibodies bound bivalently or monovalently (Fig 3 f), whose relative values become inverted when a change in concentration triggered a transition from a bivalently dominated to a monovalently dominated regime.

**Fig. 3.**
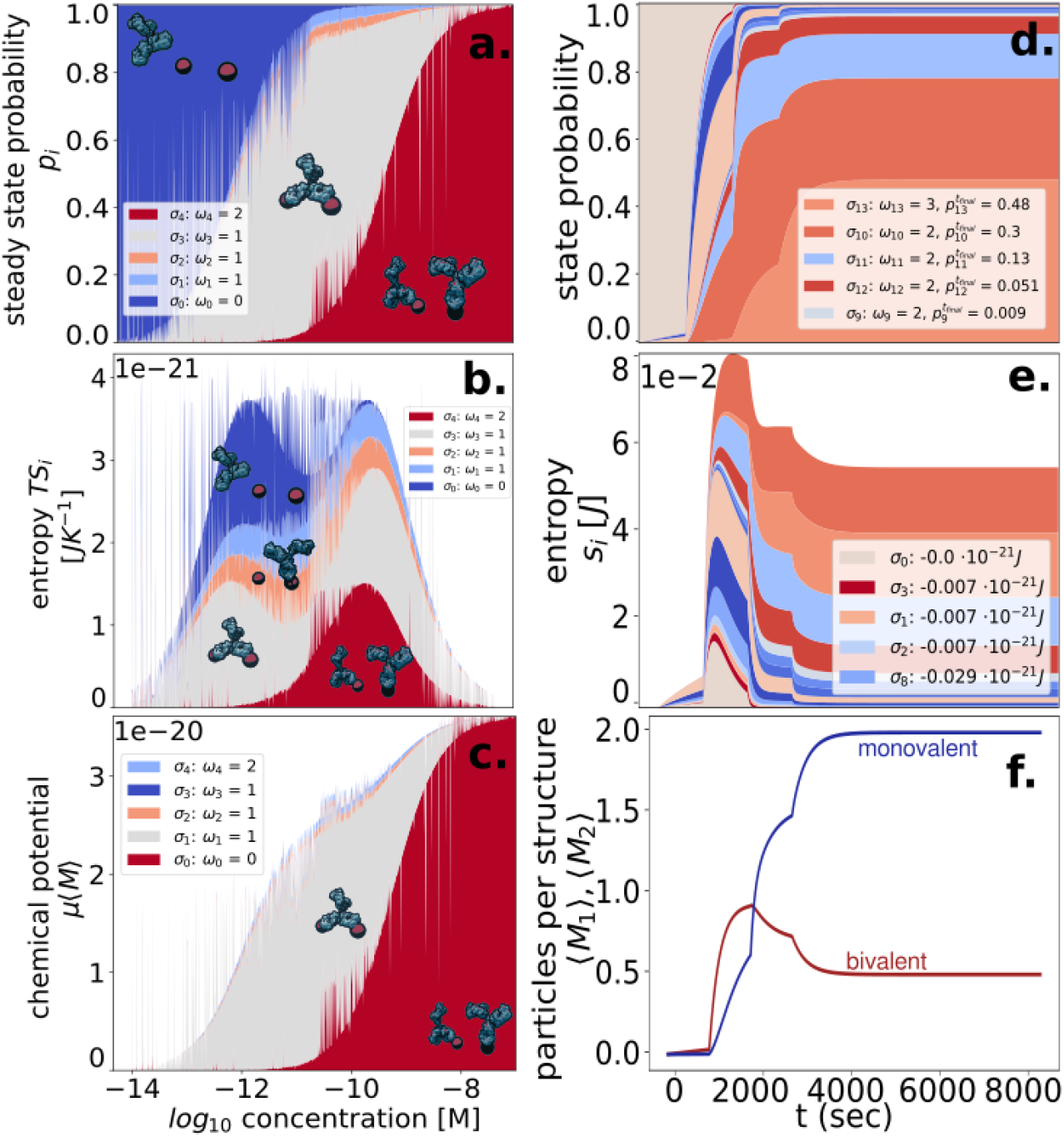
*De novo* simulation with parameterized kinetics. a: Stationary distributions colored by state of a 2-antigen system (14 nm separation) for a range of solution phase antibody concentrations demonstrating clear regions of predominantly empty, bivalent single antibody occupancy, and monovalent two antibody occupancy regimes connected by smooth transition regions. b: Distribution of state entropic contributions to free energy for a range of solution phase antibody concentrations. c: Chemical potential contributions at equilibrium from each state for a range of concentrations. (legend shows top 5 most abundant states) d: Transient state probability distribution for a simulation of a trimeric 12 × 16 × 20 nm antigen configuration with step changes in concentration. (legend shows top 5 most abundant states) e: The evolution of entropy for the trimeric system stratified by state. f: The transient evolution of monovalent and bivalent contributions of antibodies to the average number of antibodies per structure in the trimeric system. The cross between the two lines demonstrates transition between regimes by changing the concentration.

For larger systems, complete enumeration of states scales poorly with larger numbers of adjacent antigens. We developed a Markov Chain Monte Carlo implementation of the model to sample trajectories that converges to state probabilities with large sample numbers. Rather than enumerating all system states (i.e. combinations of antibodies and binding modes on a structure and possible transitions), the system performs a random walk through the large state space, computing at any point in time its rate of escape into neighboring states. We then examined collections of individual trajectories for such systems to understand their average behavior. Specifically, we examined the role of repetitive antigen spacing in simple 1D arrays. Antigens arranged according to a gradient in spacing spanning that of the region of steepest slope in Equation 1 in the range of 10 - 22 nm separation distances elicit asymmetric accumulation toward the narrow spaced end of the array (Fig 4 a), individual walking trajectories that tend toward the narrow-spaced end (Fig 4 b), asymmetric velocity (Fig 4 c), and asymmetric net displacement (Fig 4 d) according to the direction of the gradient. Antibodies binding to 1D arrays of uniform spacing exhibited divergent residence times, with antibodies spending less cumulative time on 22 nm spaced arrays (Fig 4 e) than those of narrow 10 nm spaced arrays (Fig 4 f). A comparison of net displacement also shows that antibodies moved further from their initial binding location on widely spaced arrays relative to narrowly spaced (Fig 4 g).

**Fig. 4.**
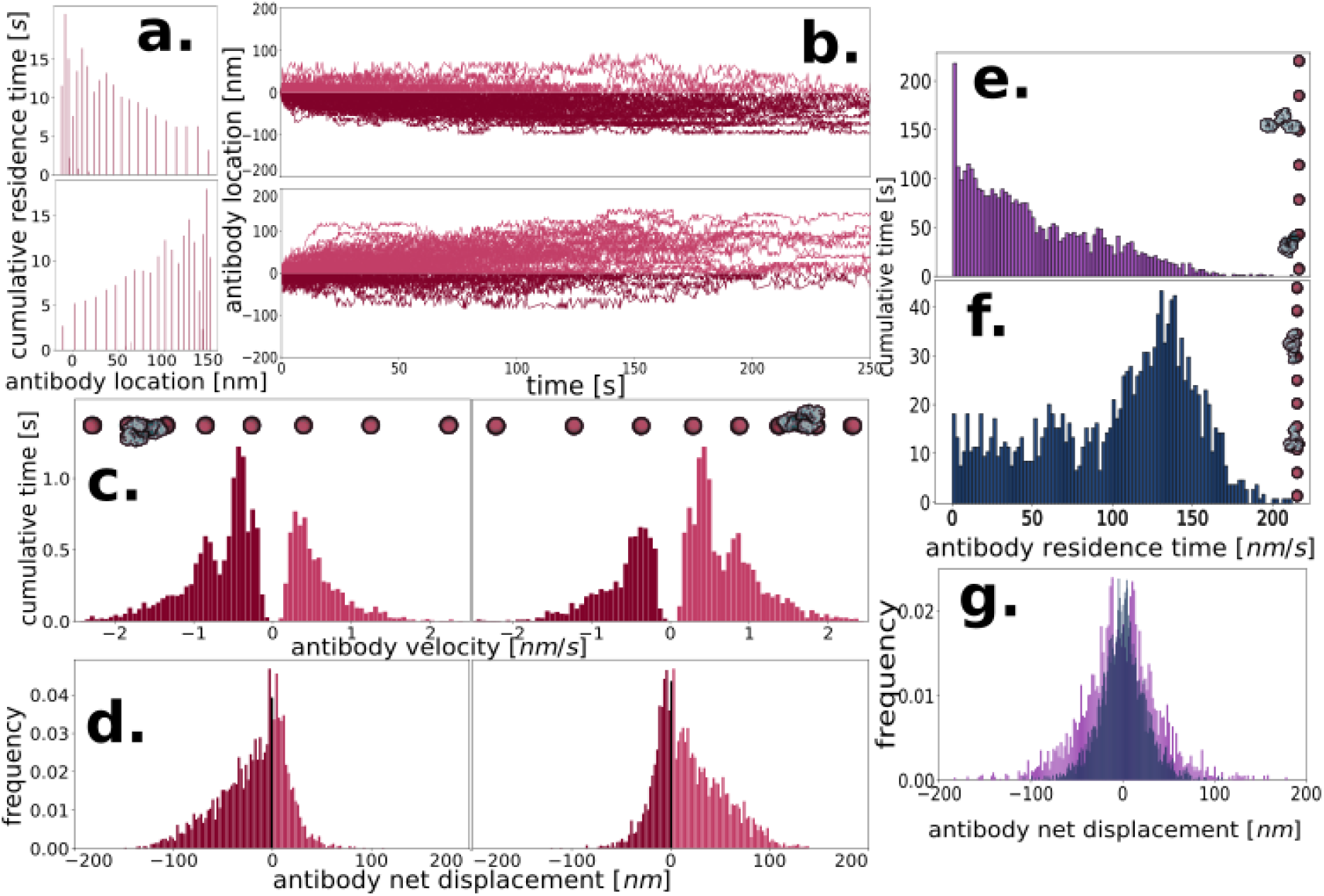
Manipulation of antibody movement through choice of antigen pattern geometry. a: Cumulative antibody residence times as a function of antibody location on 1D antigen gradients oriented with increasing spacing (top) and decreasing spacing (bottom) b: Random walk trajectories of antibodies tracked from their initial landing locations on 1D antigen gradients with increasing (top) and decreasing (bottom) spacing gradients. c: Histograms of antibody velocity on 1D antigen gradients of increasing (left) and decreasing (right) spacing distance. d: Net displacement of antibodies tracked from their initial binding location on 1D antigen gradients of increasing (left) and decreasing (right) spacing gradients. e: Histogram of antibody residence times on uniform 1D antigen array with wide 22 nm spacing. f: Histogram of antibody residence times on uniform 1D antigen array with narrow 10 nm spacing. g: Histogram of antibody displacements tracked from their initial binding locations for uniform 1D antigen arrays with wide (magenta) and narrow (blue) spacings.

## 3 Conclusion

In summary, we developed a pipeline for converting a minimal dataset from 2 PSPR experiments into a flexible stochastic model of antibody binding and walking. The model enabled us to determine steady regimes of bivalent and monovalent-dominated states depending on the solution phase concentration. We developed an enhancement to the model in order to simulate arbitrary geometries with differing separation distances between antigens by determining the binding behavior as a function of separation distance from PSPR experiments with varied spacings. This enabled us to develop an analytical expression for the change in interconversion rate as a function of antigen separation that could be applied to arbitrary geometries. The model shows that antigen patterns can influence the global movements of antibodies, with gradients in antigen spacing leading to directed migration as well as residence time control through change in uniform spacing of patterns. Taken together, this work reveals the importance of antigen pattern geometry as a control parameter in the dynamics of antibody binding and migration on multi-antigen substrates and provides a framework for modeling and exploring such phenomena using PSPR-parameterized Markov models.

## 4 Supplementary Data

### 4.1 List of model assumptions, constraints

Assume a fixed amount of bound structures that does not change with time. i.e.

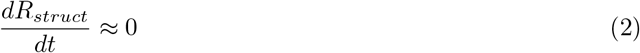

and

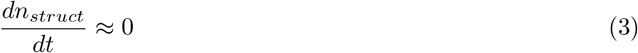

The system has an IgG reservoir that is large compared to the available binding surface and thus has an effectively fixed concentration, i.e.

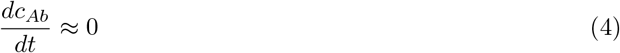

### 4.2 Conversion from SPR signal *R*_*ab*_ to bound antibody *n*_*ab*_

Consider first the simple 1-1 interaction of an antibody analyte that binds and unbinds to a structure containing a single antigen ligand.

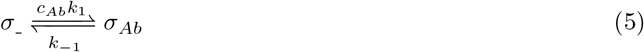

We may work in terms of molar quantities rather than concentrations or surface densities, as the dimensions of the system do not change

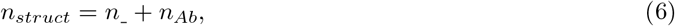

or the molar amount of structures both occupied and unoccupied [*mol*], where *n*_*Ab*_ is the number of bound antibody-structure complexes and *n*___ is the number of unoccupied structures and both where *n*_*Ab*_ and *n*___ are functions of *t*.

State probabilities are therefore:

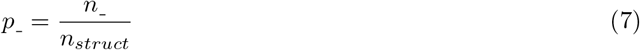

and

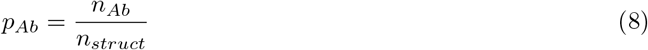

We define also an occupancy, the number of antibodies that are bound to a single structure for a given state. For simple, 1 antigen structures, this value is zero for the empty state and 1 for the bound state or

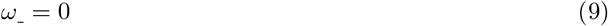

and

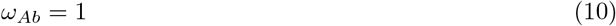

The average occupancy is a macroscopic description of the state of the system comprising *N* states, or the average fraction of bound antibodies per structure.

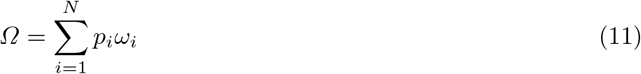

For the case of a 1 antigen structure system, this becomes

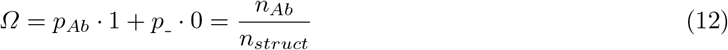

In a 1-1 binding model, change in SPR signal is proportional to the amount of bound material or in other words the change in molar amount of structures with occpupied binding sites *n*_*Ab*_.

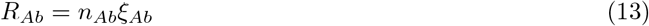

The rate of change of occupied sites is equal to the rate of conversion of unoccupied sites via binding events minus the rate of conversion of occupied sites via unbinding events.

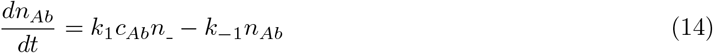

The SPR signal after the structure binding step is proportional to the molar of amount of bound structure

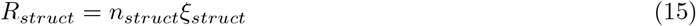

Substitute RU-based expressions of molar amounts into Equation 6:

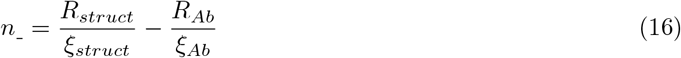

Substituting RU-based expressions of molar amounts into Equation 14 yields

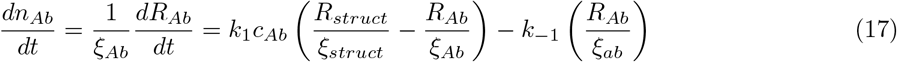

which simplifies to

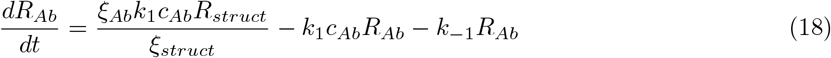

Since we have gathered both conversion constants into one term in Equation 18, we define now the occupancy signal factor

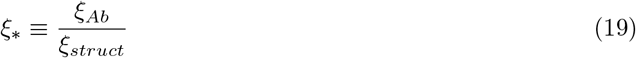

or the dimensionless ratio of molar conversion factors: bound-antibody relative to structure.

Note by rearrangement the relationship to average occupancy - i.e. the occupancy signal factor is the ratio of occupancy in terms of SPR signal to that of molar quantities.

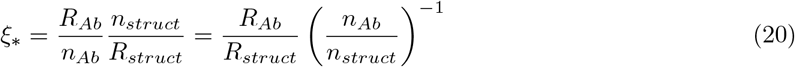

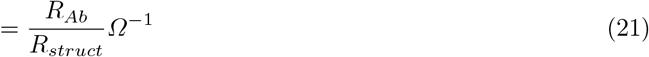

Substituting *ξ*_∗_ we then arrive at the expression for the rate of change in SPR signal with respect to time as a function of structure-binding signal and antibody-binding signal:

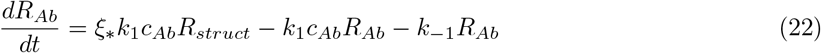

In the case of a monovalent structure (1 antigen available for binding) at the point of maximal saturation when average occupancy is unitary (*Ω* = 1), the molar quantities of bound antibody and structures are equal:

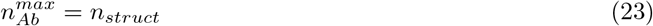

Under maximal saturation conditions, the monovalent occupancy signal factor then reduces to

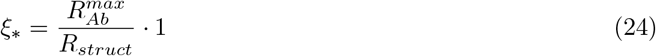

This relationship is then used to produce a standard curve from monovalent structure binding data in order to obtain the linear relationship:

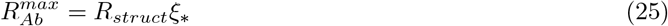

where an empirically determined *ξ*_∗_ enables one to estimate the SPR signal corresponding to an occupancy of 1 antibody per structure from the *R*_*struct*_ signal. This is useful for structures with valency greater than 1 and whose binding kinetics do not obey simple 1-1 equations. Since 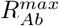 on a multivalent structure will not resemble that of the monovalent 1-1 system, we refer to this conversion factor obtained from the monovalent 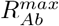 as 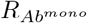, i.e. an SPR signal to antibody number conversion factor:

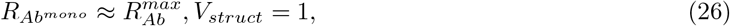

where *V*_*struct*_ is the valency.

In such cases, we obtain the average occupancy using the estimated 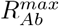 from the linear regression.

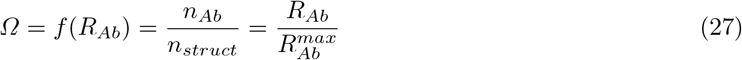

### 4.3 Empirical estimation of 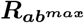

Thus, we obtain a standard curve used to convert the SPR signals for arbitrary structure configurations by empirically determining the correlation between structure binding signals and the max signals corresponding to saturated monovalent (1 antigen) structures, enabling conversion from SPR signal to occupancy in the absence of a well-understood binding model provided knowledge of the structure binding signal.

The structure bound signal (Fig 4.3 a) is taken to be the difference between signals before and after structures are flowed over the chip and allowed to bind.

Guess values of the parameters *k*_1_, *k*_−1_, and *ξ*_∗_ are supplied to a numerical minimization of the autocorrelation of residuals between experimental and theoretical curves for the 4th order Runga Kutta approximation of Equation 22, i.e. the function 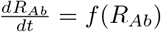 recursively approximated according to the formula

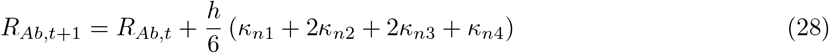

where *h* is a small timestep and the constituent terms have the form

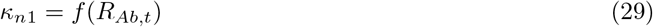

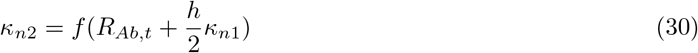

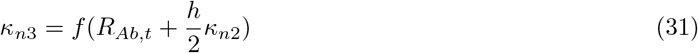

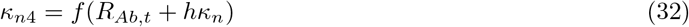

**Supplementary Figure 1.**
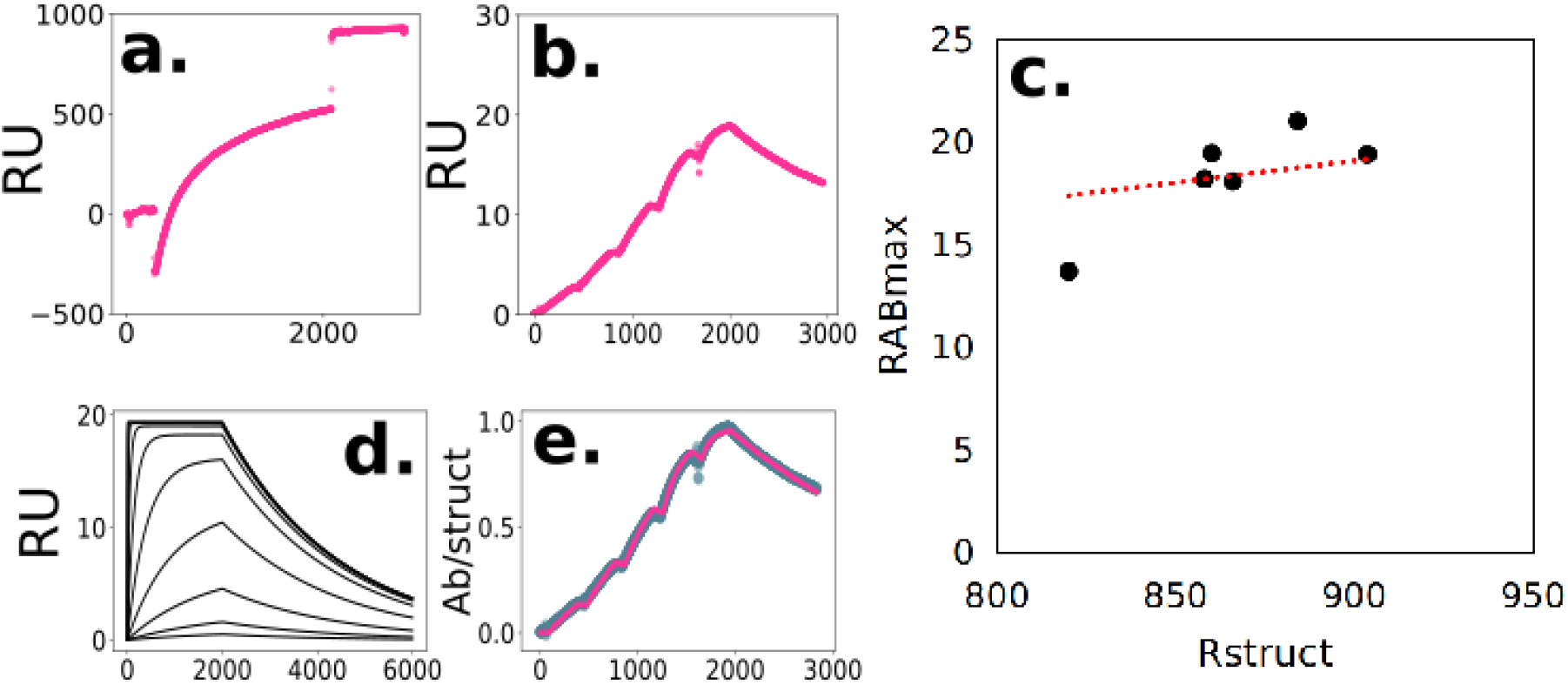
a: Sensorgram of DNA origami nanostructure binding to chip in SPR response units (RU). b: Sensorgram of antibodies binding to a 1 antigen structure proceeding the structure-binding phase in (a) shown in SPR response units (RU). c: Calculated monovalent 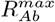 representing the number of RU corresponding to 1 antibody bound to every structure for various instances of monovalent ODE model fits to experimental binding data plotted against their corresponding structure binding signal *R*_*Struct*_ and the linear fit used to estimate the proportionality of the two variables. d: Demonstration of ODE model’s ability to predict 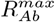, a point of saturation that the monovalent system converges to as higher concentrations push more structures into the monovalently-bound state. e: Experimental data (blue) and model (magenta) after normalization by 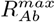, with the more useful units of antibodies per structure (AB/struct) instead of RU.

For each monovalent run (Fig 4.3 b) with a unique value of *R*_*struct*_, a projected value of 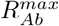 is computed with Equation 25. This enables us to make a standard curve to adjust 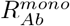 according to *R*_*struct*_ in the absence of a 1-1 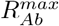 (Fig 4.3 c). This is possible because parameterization of the monovalent models by determining their rate constants enables computational prediction of 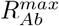 (Fig 4.3 d) in the absence of experimental saturation. We use this value as a conversion factor, enabling us to convert SPR response units into the number of antibodies per structure (Fig 4.3 e). By knowing *R*_*struct*_, we can estimate this conversion factor for non-trivial antigen configurations where the multivalence influences the ease of reaching a saturation value corresponding to 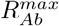.

### 4.4 Equilibrium characterization with dissociation constants

The equilibrium dissociation constant concisely describes the relationship between analyte and ligand, and provides a good basis for comparison bewteen systems across experimental conditions in which dynamic behavior can vary significantly. Given a model of the process, we can derive a formula for the equilibrium dissociation constant by solving the system of equations. For a 1-1 process we have

At steady state:

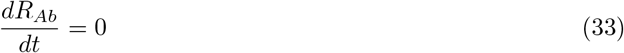

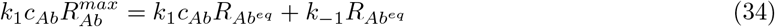

rearranging yields

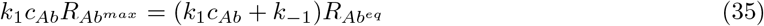

and

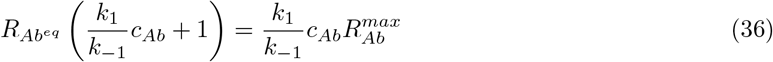

For a 1-1 monovalent model, the dissociation constant is

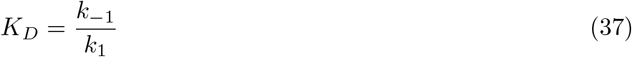

thus, at equilibrium, the SPR signal is

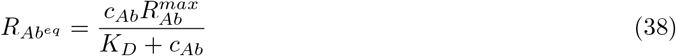

Empirical measurement of the dissociation constant is obtained by determining the equilibrium binding signals at multiple concentrations and fitting the linearized form of Equation 38 or

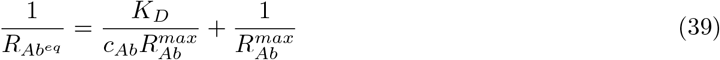

The equilibrium dissociation constant is a good descriptive parameter which captures the essential dynamics concisely.

From the dissociation constant, we know the occupancy: and the corresponding occupancy is

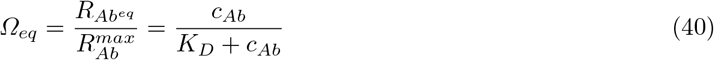

Such a concise description is desireable for complex structures as well. However difficulty arises in the case of multivalent structures which no longer exhibit simple 1-1 dynamics. One approach is to simply approximate the dynamics with a 1-1 model and obtain an apparent dissociation constant.

For the only modestly more complicated bivalent system, we can derive the relationship between an apparent dissociation constant and a complete model with two dissociation constants to describe the multiple processes taking place.

In the case of the 2-antigen structure, there are *N* = 5 total states, corresponding to an empty structure (*σ*___), two states with 1 monovalently occupied antigen each (*σ*_*Ab*__ and *σ* __*Ab*_), one state with both antigens bivalently occupied by one antibody (*σ*_.*Ab*._), and a state with both antigens monovalently occupied by antibodies (*σ*_*AbAb*_).

First, the reaction system can be represented according to the diagram in Figure 1 from the main text or the set of reactions below:

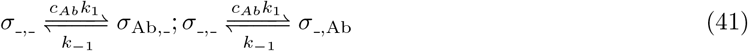

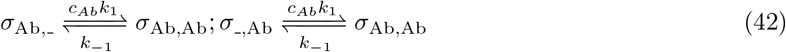

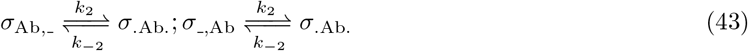

We have two dissociation constants respectively for the processes of monovalent binding and bivalent interconversion.

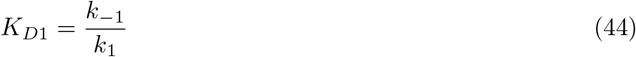

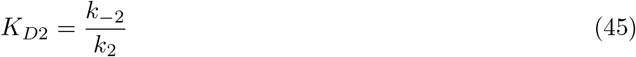

The system can be represented with a system of differential equations:

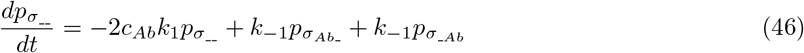

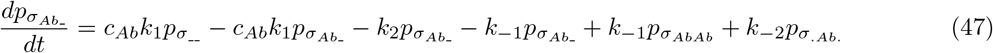

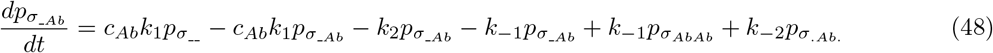

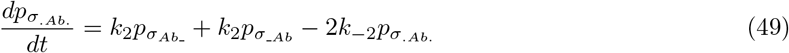

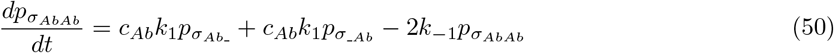

where *p*_σ__ *p*_σ*Ab* __ *p*_σ_*Ab*_ *p*_σ.*Ab*._ *p*_σ*AbAb*_ the probabilities of each of the five states in the bivalent systems, subject to the normalization condition:

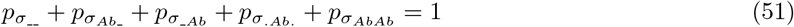

Given knowledge of the constituent equilibrium constants, we can in the simple case of the bivalent system, solve for the apparent dissociation constant as a function of the microconstants. This is, in effect, specifying a certain equilbrium value predicted on the basis of the complete bivalent model, and assuming instead that it is the result of 1-1 kinetics. However for multiple concentrations, the equilibrium will not shift proportionately, thus the apparent binding constant is a function of the concentration from which the equilibrium value is derived, making its value depent on the conditions rather than serving as a concise description of the system as a whole.

The bivalent system has, at equilibrium, the condition that the rate of change of each of its states is zero

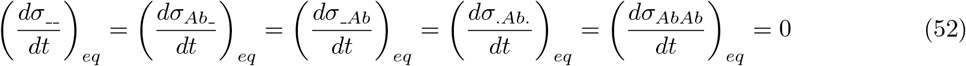

This condition plus the normalization conditions allows us to solve for the equilibrium concentrations of each of the species in terms of rate constants and the fixed solution concentration of analyte antibody.

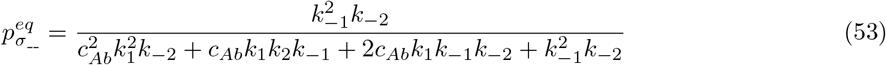

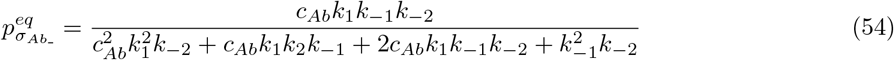

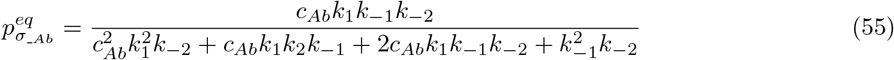

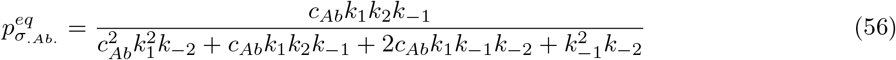

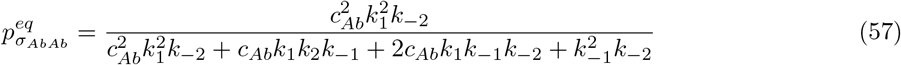

We can combine states according to their correponding occupancy - i.e. the number of antibodies that the state contributes to the overall signal due to bound antibody, where

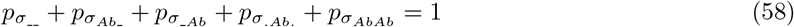

The probabilistic definition of occupancy is the expectation value of state occupancy. Each state has a corresponding integer occupancy associated with the number of antibodies bound to the structure in that state as well as a respective probability of that state at any point in time. The equilibrium occupancy is thus the average occupancy of all the states weighted by their equilibrium probabilities.

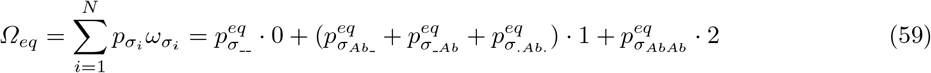

substituting Equations 53 through 57, we arrive at

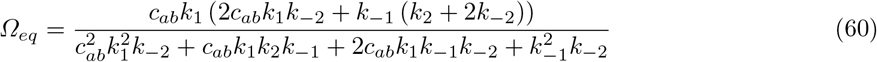

### 4.5 Apparent dissociation constant

Taking the 2-antigen system equilibrium occupancy from Equation 60 and applying it to the equilibrium occupancy in terms of the 1-1 dissociation constant Equation 40 can be used to solve for an apparent equilibrium dissociation constant of the form

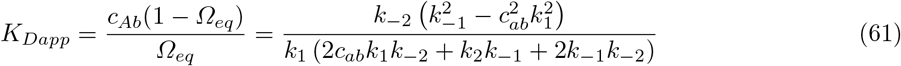

This constant is a value that would be obtained from a 1-1 fit to an equilibrium SPR value that arose from the 2-antigen kinetics. Rearranging and substituting Equations 44 and 45 into Equation 61, the formula simplifies to

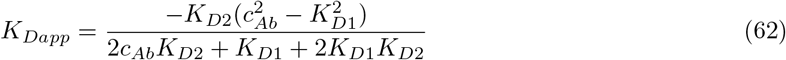

which we may note is a function of concentration, and which has a root at the critical value when 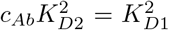, i.e. the point at which average equilibrium occupancy greater than 1 is expected in the 2-antigen system, and rendering impossible any 1-1 kinetic description.

Rearrangement of Equation 62 enables us to determine the interconversion constant from an apparent dissociation constant provided that we know the monovalent binding constant.

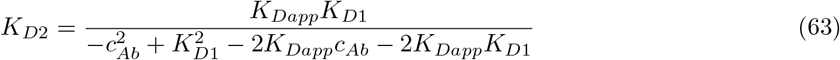

**Supplementary Figure 2.**
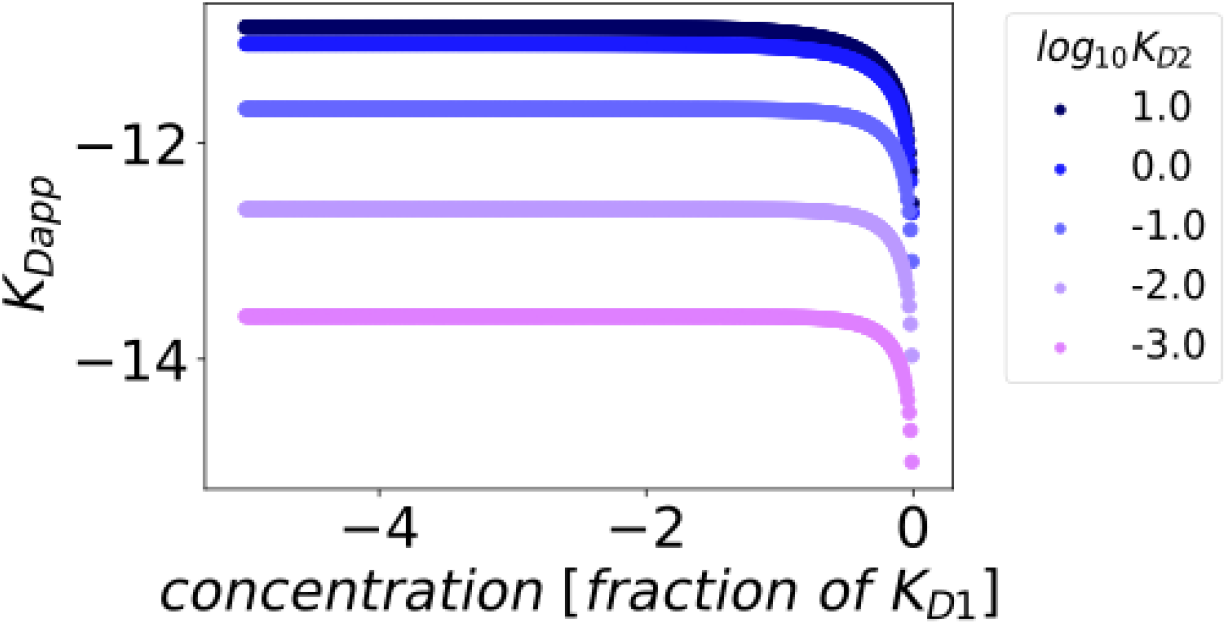
Log-log (base 10) plot of the apparent dissociation constant as a function of concentration represented as a fraction of the monovalent dissociation constant. For concentrations approaching that of the monovalent dissociation constant, the apparent binding constant changes significantly, whereas low concentrations lead to less concentration dependence.

At concentrations where *c*_*Ab*_ ≪ *K*_*D*1_, the relationship between 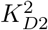 and 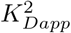 is relatively constant (Fig 2). Note that this is only valid for PSPR data with a 2-antigen topology of a single separation distance.

### 4.6 Mathematical description of spatial tolerance

Spatial tolerance refers the favorability of bivalent antibody binding according to the spatial distribution of the 2 adjoining antigens. Some antibodies stretch and compress more than others leading to a greater chance of entering and remaining in a bivalent state. In our model, we propose that the monovalent binding step occurs separately from the bivalent binding step, and that it is purely dependent on the solution phase concentration and the epitope-paratope binding affinity. Spatial tolerance therefore is a property of the interconversion step from monovalent to bivalent states and the reverse process from bivalent back to monovalent. For antigens separated by very small distances, electrostatic repulsion in response to compression and steric hindrance within the IgG molecule occurs, penalizing conversion to bivalent binding and/or favoring unbinding back to monovalent states. Conversely, at larger separation distances, the molecule must stretch to accomodate the gap, again penalizing conversion to bivalence and/or favoring conversion back to monovalence.

Spatial tolerance is a description of the landscape of this tradeoff - the breadth of the favorable region in between extremes that is conducive to bivalent binding, the sharpness and degree of symmetry of the transitions to monovalent preference at close and far separations. We can model spatial tolerance phenomenologically with an equation for determining the interconversion constant *K*_*D*2_ as a function of the separation distance between two antigens *x*:

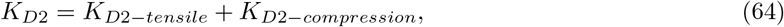

where *K*_*D*2−*compression*_ (Fig 3 a) and *K*_*D*2−*tensile*_ (Fig 3 b) are respectively exponential and logistic terms. These model separately the decrease in interconversion due to tensile stretch of the molecule at increasing distances and that due to the onset of exluded volume, electrostatic repulsion, or steric hindrance caused by compression of the molecule to bridge close distances.

**Supplementary Figure 3.**
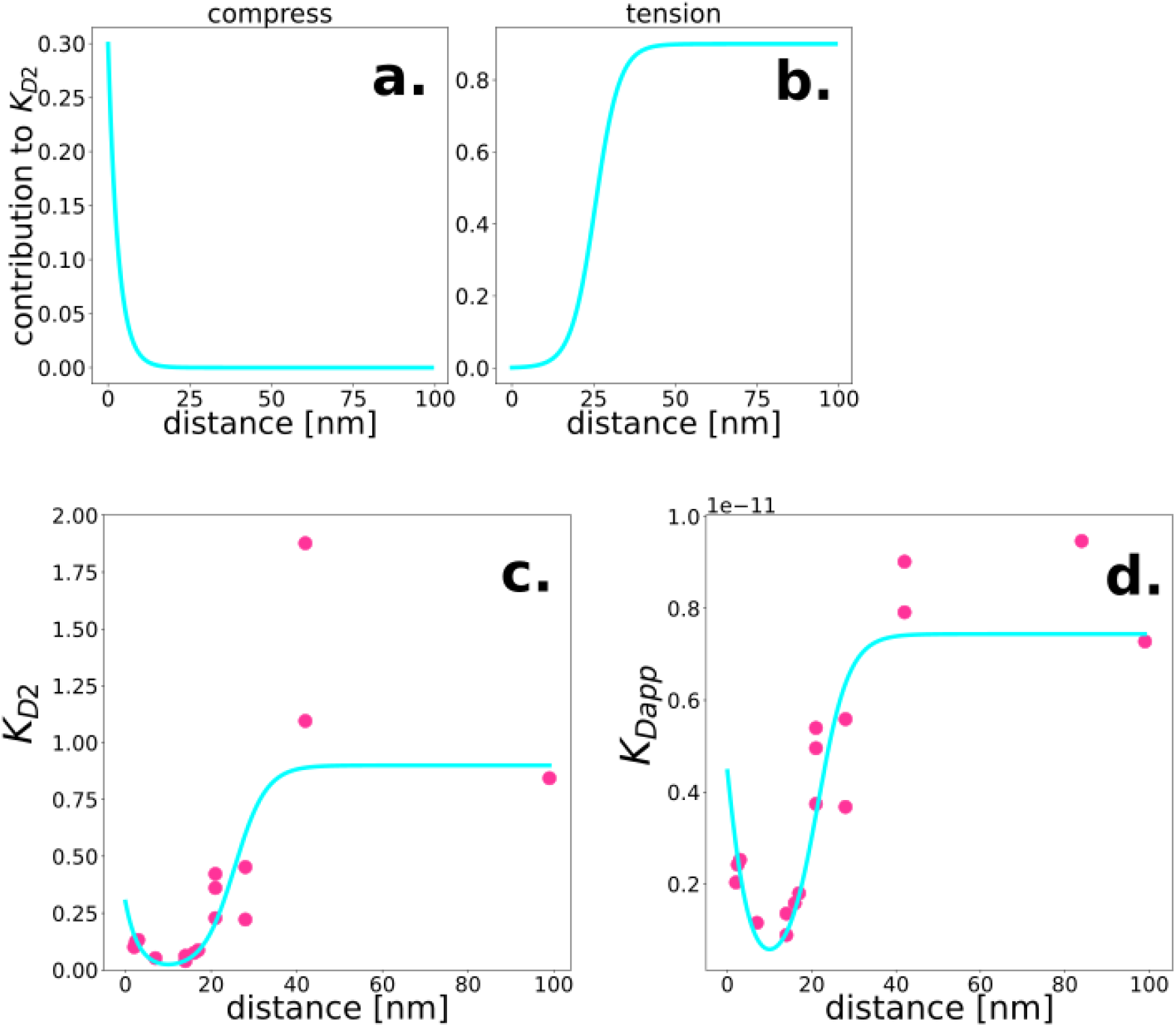
Measurment and modeling of spatial tolerance of antibody with increasing spatial separation of 2 antigens. a: Exponential compression term of spatial tolerance plotted alone versus distance. b: Sigmoidal tension term of spatial tolerance plotted alone versus distance. c: The spatial tolerance function, experimental interconversion constant values (magenta) and the fitted spatial tolerance model (cyan) both plotted against separation distance of two antigens. d: The apparent dissociation constant assuming 1-1 binding kinetics for what is actually a 2-antigen bivalent system calculated from interconversion constant data on the basis of equilibrium occupancy predicted by the bivalent model in terms of individual rate parameters.

The tensile term is built from a logistic function and has the form:

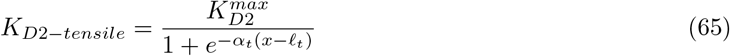

where 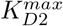 is an upper limit of the value of *K*_*D*2_, *α*_*t*_ is the logistic growth rate or steepness with which the tensile penalty grows at increasing separation distances and has units of inverse length, and *ℓ*_*t*_ is the value of the midpoint of the sigmoidal curve which can be thought of as a characteristic length that defines the scale below which favorable interconversion will occur and above which the function approaches minimal interconversion.

The exponential compressive term has the form:

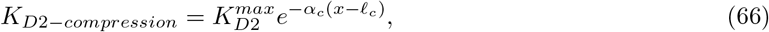

where *α*_*c*_ is the exponential decay rate which has units of inverse length, and *ℓ*_*c*_ is another characteristic length parameter with units of length. The model is subject to the constraint *ℓ*_*c*_ < *ℓ*_*t*_.

The combined expression yields Equation 1 which predicts the interconversion constant as a function of separation distance (Fig 3 c). This can be converted to an effective or apparent dissociation constant as if a 1-1 model on the basis of the bivalent model’s prediction of equilibrium occupancy (Fig 3 d) - see Section 4.5.

### 4.7 Markov model of arbitrary antigen pattern geometries

For the binding kinetics of multi-antigen patterns of systems of sizes on the order of 2-8 adjacent antigens, we employ a fully enumerative Markov chain model based on a complete transfer matrix, i.e. all possible states and transitions of the system. The antigen pattern itself is modeled as a discrete network of antigen sites with a Euclidean distance matrix

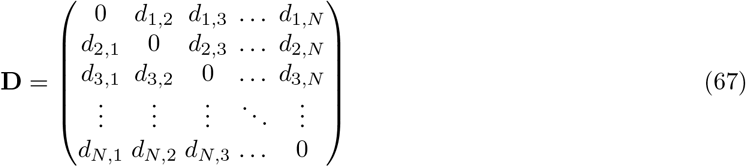

An adjacency matrix is formed by applying a critical distance cutoff to **D**, such that long range interactions will be ignored, greatly reducing the number of states and transitions to enumerate.

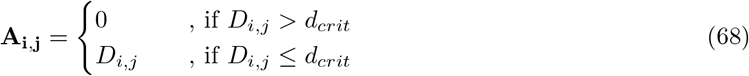

A single state *σ*_*i*_ of the system is defined as a set of antigens, their status (empty, monovalently occupied, bivalently occupied) and a pointer indicating of which bivalent-status antigens are linked to each other. The state space of a system is the set of all states that a structure in the system can assume, i.e. 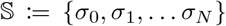. The set of states are thus all the possible configurations of empty, monovalently bound, and bivalently bound antibodies given the constraints of the pattern geometry (Fig 4).

Each state is linked to adjacent states by elementary transitions, i.e. the change in status of individual antibodies comprising the state. Those transitions are either the concentration-dependent addition or the subtraction of a single antibody to the system via monovalent binding or unbinding:

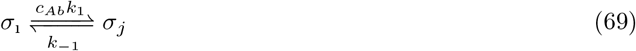

or a bivalent interconversion event where a monovalently bound antibody binds to an adjacent antigen site, changing its status to bivalently bound and vice versa:

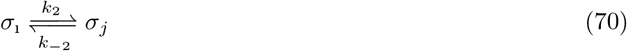

**Supplementary Figure 4.**
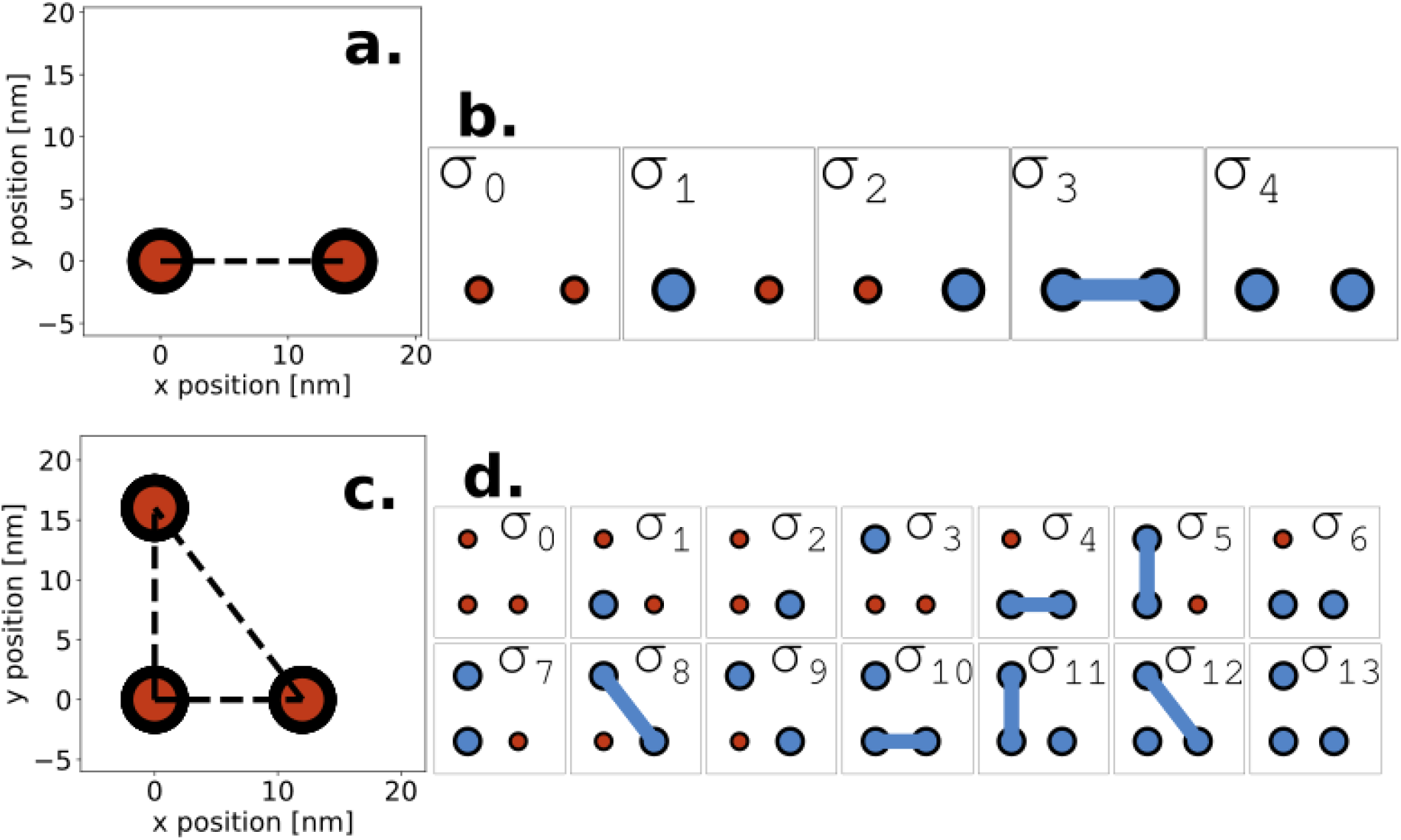
Enumerated states for basic and arbitrary geometries. a: Dimensions of a basic bivalent antigen pattern with 15 nm separation between antigens. b: The set of possible states for the basic bivalent geometry with blue dots indicating monovalently bound antibodies, blue lines connecting two dots indicating a bivalently bound antibody, and red dots indicating empty antigen sites. c: The dimensions of an arbitrary geometry, a 12 × 16 × 20 nm right triangle. d: The set of states corresponding to the right triangle geometry.

The system parameters are the set of zero order transition rates {*r*_1_ = *c*_*Ab*_*k*_1_, *r*_−1_ = *k*_−1_, *r*_2_ = *k*_2_, *r*_−2_ = *k*_−2_. The multi-antigen-antibody system is thus fully described by the continuous time Markov model 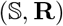 defined as its set of states and its corresponding transition rate matrix of the form:

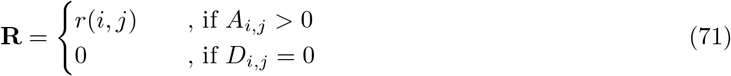

Automated enumeration of states in systems of arbitrary antigen pattern geometry is accomplished using an implementation of the breadth-first search (BFS) algorithm. The algorithm searches for edges between adjacent states and assigns the appropriate elementary rate process. A queue of neighboring states is made upon the visitation of any state. One-by-one, the algorithm visits each state in the queue, populating it with additional states when they are discovered, and skipping the addition of states that have already been visited. The algorithm thus is characterized by an initial expansion phase of the queue followed by a systematic reduction of the queue until all states have been visited, and the queue becomes empty. This exhaustive enumeration is deterministic, and enables us to assemble a complete transition matrix regardless of antigen geometry. However as the number of adjacent antigens grows, the number of combinations increases dramatically, thus for larger systems, a sampling based approach must be used instead.

### 4.8 Transient (non-equilibrium) dynamics of enumerative PSPR models

The continuous time Markov model enables us to compute transient evolution of the system. The probability distribution

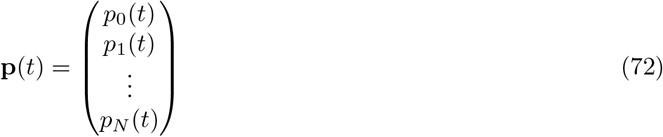

is a vector whose elements *p*_*i*_(*t*) are the probabilities of the respective system states *σ*_0_, *σ*_1_, … *σ*_*N*_ at time *t*. A uniform probability distribution would, for example, represent equal probabilities of finding a structure in any one of the possible states. Or another example is at the start of a single cycle kinetics PSPR run, when the initial condition **p**(*t*_0_) is that of a distribution where *p*___(*t*_0_) = 1 for the state *σ*___ corresponding to an empty structure and *p_i_*(*t*_0_) = 0 for all other states.

The transient evolution of state probabilities is computed from an intitial condition using the linear system of Chapman-Kolmogorov differential equations:

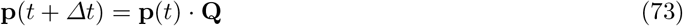

making use of an infinitessimal generator matrix **Q** which is obtained from the rate matrix and used to determine the relative rates at which state probabilities change with incremental time.

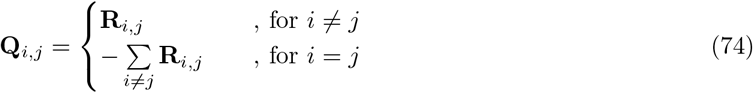

The infinitessimal generator is then used to compute the change in state probability distribution going from one time point to the next by the matrix exponential formula:

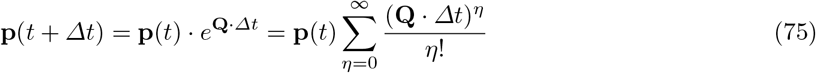

where *η* is the computation’s depth of recursion - the higher the more accurate, and *Δt* is an incremental advancement in time. Due to numerical instability of this solution, we employ the uniformized discrete time Markov chain method of Fox and Glynn in order to stably compute Equation 75 [4]. The continuous Markov model 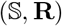 is approximated by a discrete model 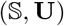 by renormalizing the generator matrix with respect to the fastest outgoing rate or the uniformization rate *q*:

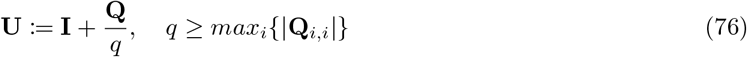

where **I** is the identity matrix. Equation 75 becomes the approximation

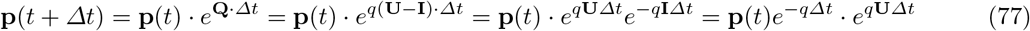

The matrix exponential is then approximated with the following Taylor series expansion:

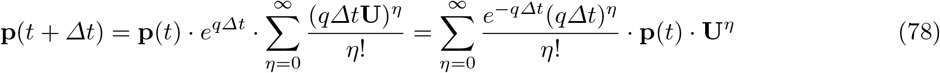

Using Equation 78, we can stably compute the transient evolution of a system from an initial condition. The system entropy can by computed by

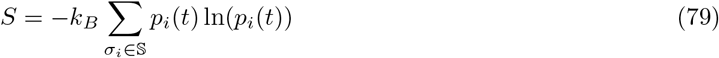

### 4.9 Fitting continuous time Markov models to PSPR data using autocorrelation of residuals

Using Equation 78 to compute the transient probability distribution of the system, we are able to also compute the occupancy at each time point using the definition from Equation 11. The system occupancy is thus a function of time of the form

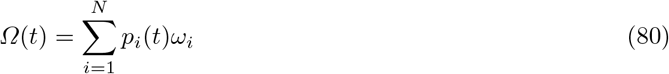

The continuous time Markov model is fitted to experimental data by comparing occupancies computed on the basis of Equations 78 and 80 with that of occupancy computed from normalizing PSPR data via Equation 27 and we can see that the theoretical curve either correctly or incorrectly fits the experimental data depending on the parameterization (Fig 5 a and Fig 6 a). Residuals (Fig 5 b and Fig 6 b) are computed by

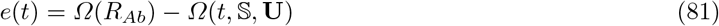

While fitting by minimizing the sum of squared residuals can be used to obtain acceptable model parameterizations, we used residual analysis with autocorrelation to improve the robustness of fitting and reduce systematic mis-parameterization by making fits more sensitive to divergence in curve shapes. We compute an absolute, average autocorrelation over a fixed interval *kΔt* with *k* = 50 by :

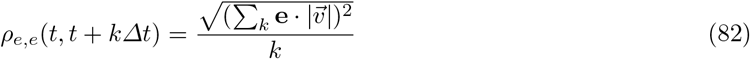

where **e**(*t, k*) = [*e*(*t*), *e*(*t* + *Δt*), … *e*(*t* + *kΔt*)], **v** = [0, 1, … *k*], and |**v**| is the conjugate of **v**. The objective function numerically minimized *min*(*ε*) to obtain fits to experimental PSPR data is then the sum squared residual vector weighted by its autocorrelation vector:

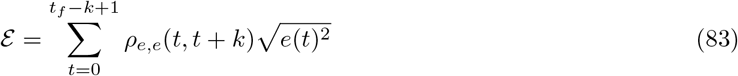

**Supplementary Figure 5.**
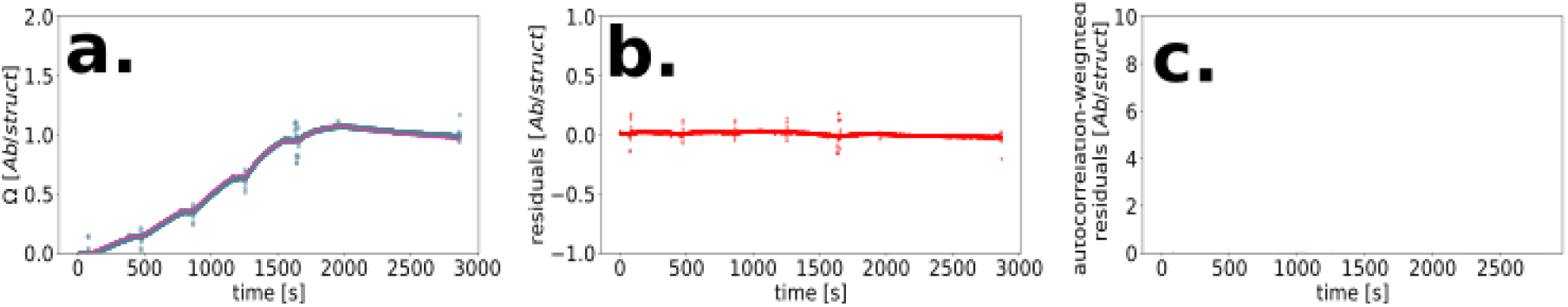
Example of a well-parameterized bivalent model, with strong agreement between experimental and theoretical occupancy. a: Markov model (magenta) prediction of average occupancy (antibodies per structure) superimposed on experimental PSPR binding data for a 2 antigen pattern (blue). b: Residuals comparing the theoretical and experimental curves in a. c: Residuals from b weighted by autocorrelation.

**Supplementary Figure 6.**
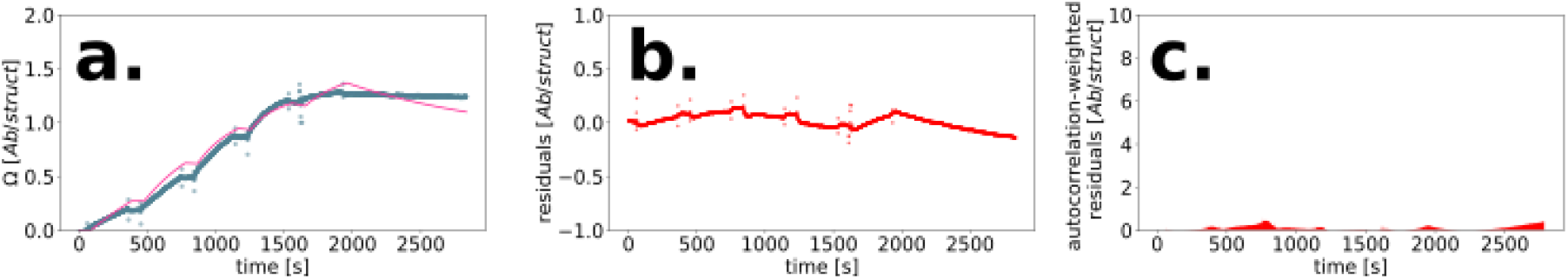
Example of a misparameterized bivalent model with visual misalignment between theoretical and experimental occupancy curves and corresponding divergence of residuals. a: Markov model (magenta) prediction of average occupancy (antibodies per structure) superimposed on experimental PSPR binding data for a 2 antigen pattern (blue). b: Residuals comparing the theoretical and experimental curves in a. c: Residuals from b weighted by autocorrelation.

This provides an error function sensitive to sustained divergence of model and experimental data (Fig 6 c) even if the two curves cross paths, and like summing the residuals provides a low value when alignment is good (Fig 5 c).

**Supplementary Figure 7.**
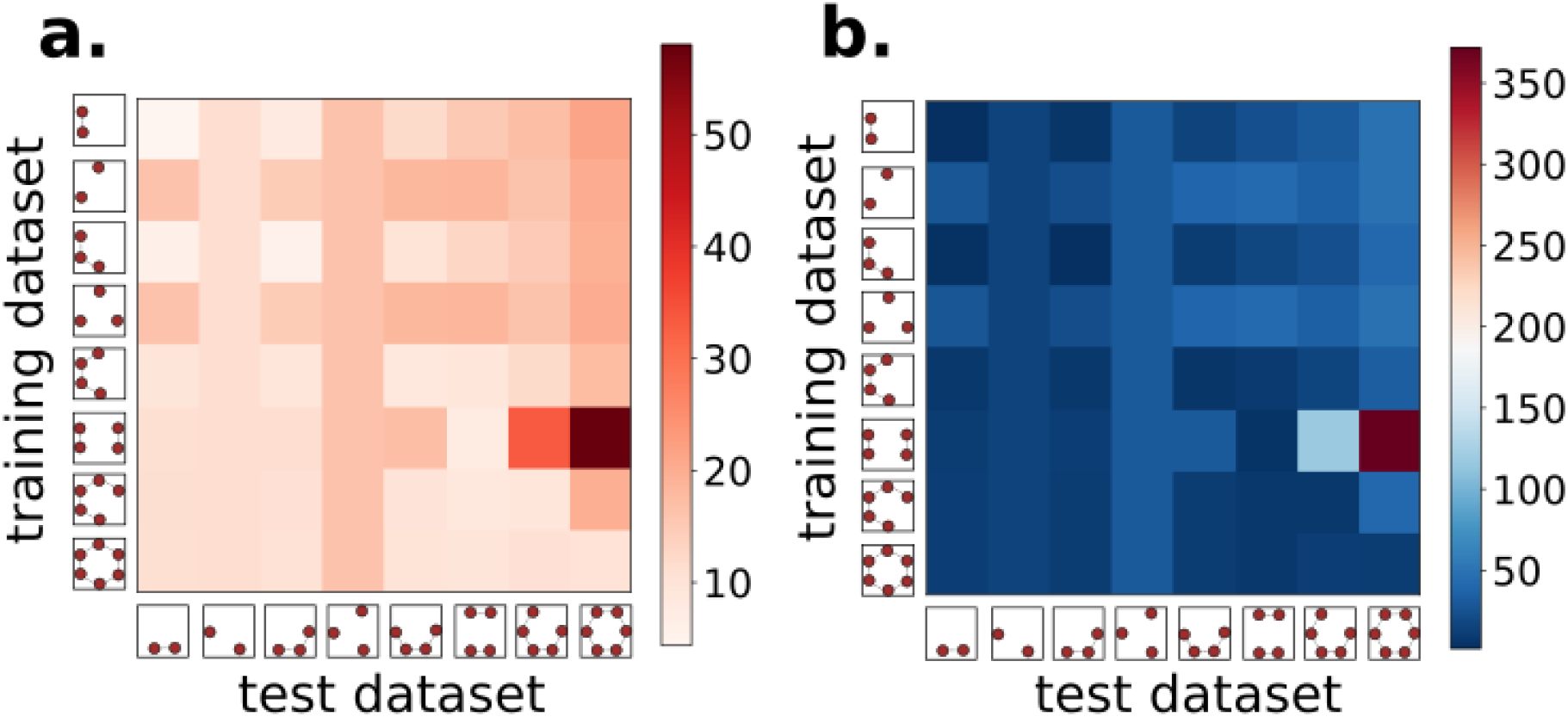
a: Matrix depicting the sum of residuals (color map) for Markov models parameterized on experimental binding data from antigen patterns on the vertical axis (training dataset) and performance-tested against other patterns (test dataset). b: Same as (a) except that colormap indicates the residuals weighted by their autocorrelations.

We performed cross validation of Markov model fitting by parameterizing based on experimental data from various antigen patterns (all with fixed nearest-neighbor separation distances between antigens to remove the complication of separation distance dependence of the binding kinetics). The rate parameters derived from these training data were then fixed and the model was applied to other patterns as a limited test of extrapolation of a parameterized model to different antigen pattern geometries. Absolute sums of residuals (Fig 7 a) and absolute sums of residuals weighted by autocorrelation (Fig 7 b) show that the best-performing models were the those that are the most complex and exhibiting bivalence such as the hexagonal and pentagonal configurations in the last two rows. This suggests that downward extrapolation in pattern complexity is more viable than upward.

### 4.10 Determination of thermodynamic properties

We can obtain equilibrium probabilities from the uniformized CTMC first by simulation out to long time scales at fixed solution concentration until probabilities cease to change:

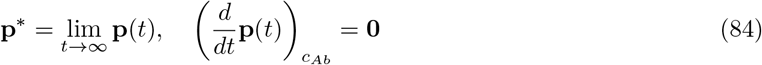

We can determine the steady state probability distribution more expediently on the basis of the infinitessimal generator matrix, Equation 74, solving numerically for the probability distribution which, when multiplied with the generator matrix produces a vector of zeros, meaning that there is zero change from one moment to the next, subject to the normalization condition whereby all probabilities must sum to 1. I.e. the steady state probability distribution is the solution to the matrix equation

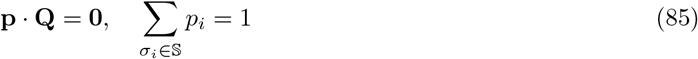

The multi-antigen structure in the context of a PSPR experiment is an open system, freely allowed to exchange particles with the large external reservoire connected to it. With (*T, V, c*_*Ab*_) held constant, the system (an antigen patterned structure) will approach a minimum free energy at steady state by exchanging antibodies with the bath, obeying the Boltzmann distribution law

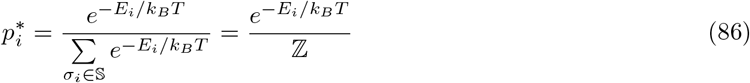

where ℤ is a grand canonical partition function which predicts equilibrium at a grand potential free energy minimum *dΦ*(*T, V, μ*_*Ab*_) = 0 with chemical potential *μ*_*Ab*_ = −*k*_*b*_*T* ln *c*_*Ab*_, and *E*_*i*_*p*_*i*_ = *μ*_*Ab*_ + *μ*_*mono*_*n*_*mono*_ + *μ*_*biv*_*n*_*biv*_ are state energies determined by the environmental potential due to solution phase antibody concentration as well as the individual potentials of antibody monovalent and bivalent bonds populating the state, *n_mono_* monovalent bonds and *n*_*biv*_ bivalent bonds with respective chemical potentials *μ*_*mono*_ and *μ*_*biv*_. The value of ℤ and the state energies are solved numerically for a given steady state probability distribution, making it possible for us to determine thermodynamic quantities.

We can obtain thermodynamic quantities such as the solution concentration dependent equilibrium grand potential free energy via:

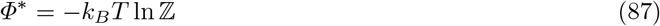

The equilibrium probabilities enable us to calculate relative potential differences via

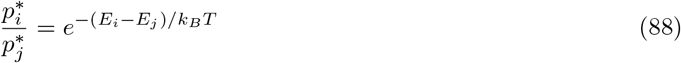

This makes it possible to compute chemical potentials of monovalent and bivalent bonds, for example from the basic 2-antigen system of a fixed separation, by comparing equilibrium probabilities in states that differ by exactly 1 bond of a given type. This is equivalent to obtaining the change in free energy via the dissociation constant for that process.

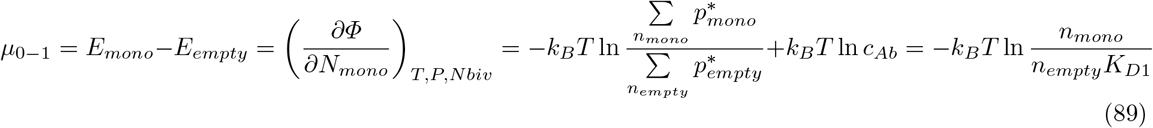

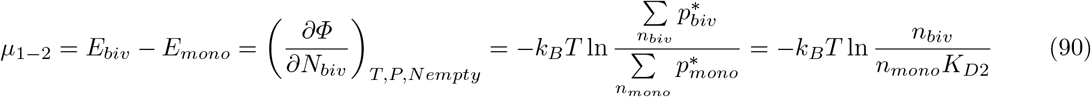

where *n*_*empty*_, *n*_*mono*_, and *n*_*biv*_ are the respective degeneracies of empty, monovalently 1-occupied, and bivalently 1-occupied states. For the 2 antigen system, these values are respectively 1, 2, and 1. For the rabbit IgG, anti-digoxygen model and a separation distance of 15 nm, the chemical potentials are *μ*_0−1_ = 1.805 × 10^−20^ **J** / particle and *μ*_1−2_ = 8.889 × 10^−21^ **J**/particle.

We can also obtain a stand-alone bivalent binding chemical potential such that talleying the number of monovalent and bivalent molecules in a state would yield the potential of that state.

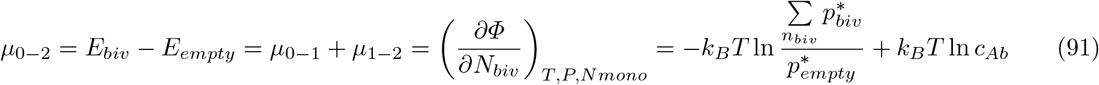

This gives us *μ*_0−2_ = 2.693 × 10^−20^ **J** / particle for the 15 nm separation. By this method, the distance-dependent *K*_*D*2_ model of Equation 1 can be converted to a chemical potential curve.

### 4.11 Markov chain Monte Carlo version of model

Systems with larger numbers of bivalent connections have many states and transitions, and the fully enumerative CTMC does not scale well. For these systems, we use a Markov Chain Monte Carlo (MCMC) sampling approach whereby only local states are computed throughout the trajectory of a single system. Multiple trajectories are sampled to approach and approximate the probability distributions that would otherwise be computed exactly for the enumerative method. Instead of computing the flux in state probabilities over fixed intervals of time, we instead computing Poisson intervals (dwell times) of states according to the rates of processes that each state is subject to.

## 5 Experimental Methods

The experimental data used in this work was presented previously as supplementary control data in [11] and was not analyzed further than a basic fitting to a standard 1-1 model. In the current work, we have used the data to develop a more accurate mechanistic model, a pipeline for constructing such models from a minimal dataset, an in-depth physical characterization framework, and *de novo* simulation that go beyond previous work.

### 5.1 Reagents

Oligonucleotides (unmodified and digoxigenin-modified) were purchased from IDT (Belgium) in 96-well plates format. Chemicals (NaCl, KCl, MgCl_2_, Tris-HCl, EDTA, PEG800, NaOH, KOAc, KOH, NaOAc) for buffer preparation were purchased from Sigma-Aldrich. Streptavidin was purchased from Sigma-Aldrich. Phosphate-buffered saline 1M stock solution was purchased from Sigma-Aldrich. BIAcore consumables (CM3 sensor chip, HBS-EP running buffer) were purchased from GE Healthcare. Amicon centrifugal filters with 100 kDa MWCO were purchased from Merk Millipore.

### 5.2 Origami design and production

The 18-helix bundle DNA origami nanotube was designed with caDNAno[**?**], using the honeycomb lattice. This structure has been characterized earlier[11, **?**,**?**,**?**]. In short: the p7560 scaffold was extracted from M13 phage, and the 18-helix bundle DNA nanotube was folded under the following condition: 20 nM scaffold, 100 nM each staple oligonucleotide, 13 mM MgCl_2_, 5 mM Tris pH 8.5, 1 mM EDTA. The mixture was subjected to heat denaturation at 80°C for 5 min followed by a slow cooling ramp from 80°C to 60°C over 20 min and 60°C to 24°C over 14 hr. The excess staples were removed by ultrafiltration with Amicon 100 kDa MWCO filters. The wash buffer used was 1× PBS supplemented with 10 mM MgCl_2_.

### 5.3 Pattern surface plasmon resonance protocol

The BIAcore t200 instrument from GE Healthcare was used for surface plasmon resonance experiments. The running buffer used in all experiments is HBS-EP supplemented with 10 mM MgCl_2_. The flowrate used for all kinetic experiments is 30 *μl/min*. Streptavidin was diluted to a final concentration of 10 *μg/ml* in 10 mM NaOAc pH 4.5 and chemically attached to the CM3 sensor chip with NHS/EDC coupling, using the standard protocol from GE Healthcare. Anchor oligonucleotides containing a 3’ biotin modification were diluted to 200 nM in 1 × HBS-EP running buffer and was injected over the surface for 20 min followed by washing of nonspecifically bound oligos by injecting 50 mM NaOH for 5 mins. The DNA nanostructures were diluted to 5 nM and injected over the surface for 20 mins followed by washing with running buffer for 10 mins. Antibodies were diluted to various concentration in running buffer ranging from 0.025 nM to 0.5 nM. The single cycle kinetics injection method was used to sequentially inject the antibody solution over the surface, started from the lowest concentration, the contact time for each concentration is 5 min. After the final antibody injection, the dissociation curve was recorded for 15 min. The immobilized DNA nanostructures and bound antibodies were removed from the surface by injecting 50 mM NaOH for 5 mins and then the surface is ready for the next round of experiment. The t200 evaluation software was used initially to fit the acquired data, for this we used a 1:1 Langmiur binding model to fit the data and estimate the *k*_*a*_, *k*_*d*_, *K*_*D*_ and antibody binding capacity.

